# Falafel-Style Wrapping of AuNPs for DNA Origami Barcodes

**DOI:** 10.64898/2026.05.18.725969

**Authors:** Sarah Youssef, Thomas Swope, Thorsten L. Schmidt, Diana P. N. Gonçalves

## Abstract

The ability to encode and reliably read nanoscale information is increasingly important for multiplexed biomolecular detection and super-resolution imaging. DNA origami provides a uniquely programmable platform for arranging structural and functional elements with nanometer precision, enabling the creation of identifiable nanoscale patterns. In this context, DNA origami-based barcodes that incorporate gold nanoparticles (AuNPs) to encode either origami geometry or the identity of specific biological targets within defined nanoparticle patterns have been paired with transmission electron microscopy imaging for decoding. However, surface-bond AuNPs may detach during handling, purification, or biological incubation, leading to misidentification or decoding errors in barcode analysis. Here we report a rational design for the controlled encapsulation of AuNPs within DNA origami tubes to enhance nanoparticle retention and structural integrity. We engineered curvature-inducing modifications in a flat rectangular DNA origami scaffold to promote inward folding and confinement of AuNPs. These barcodes can be further functionalized on the outer surface with bioactive aptamers and/or fluorescence dyes, enabling targeted interactions with cells and optical readout. Programable dimerization further expands multiplexing capacity. This design provides a robust framework for structurally stable origami barcodes and advances the development of high-resolution, multiplexed labeling and diagnostic platforms.

**Graphical abstract:** 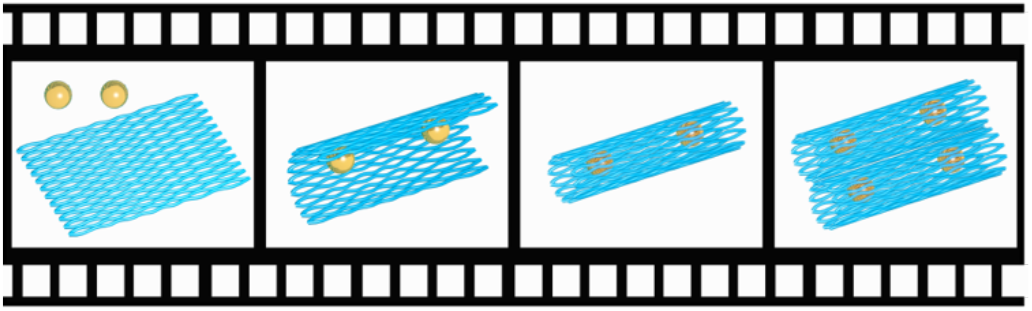

## Introduction

The increasing demand for multiplexed detection and identification of diverse biomarkers, such as proteins and nucleic acids has driven the development of several origami-based barcoding platforms to facilitate efficient sample handing and high-throughput analytical workflows.^1–7^ DNA origami is a technique in which a long single-stranded (ss) DNA scaffold is folded into predefined two- and three-dimensional (2-, 3D) architectures *via* the hybridization with approximately 200 to 250 short “staple” strands.^8–11^ Since each staple strand has a unique sequence and defined position within the structure, DNA origami provides addressable binding sites with spatial resolution on the order of approximately 3 nm. This degree of programmability enables the precise organization of diverse functional components, including metal nanoparticles, proteins, and other biomolecules.^1,12–19^ An example of the versatility and programmability of DNA nanotechnology is the development of gold nanoparticle (AuNP) based DNA origami barcodes. These constructs consist of DNA origami structures decorated with defined spatial patterns of AuNPs, which can be read out by Transmission Electron Microscopy (TEM).^1,2^ The concept behind these particular origami barcodes is to encode structural identity or the presence of specific functional elements, directly into the nanoparticle pattern on a given origami template. Recently, we have demonstrated that such barcodes can be applied for the immunolabeling of proteins in cryosectioned mouse retina tissue.^1^ However, depending on the intended application, particularly in the context of biological studies, the functionalization of the origami surface with AuNPs can introduce unintended structural and functional consequences. Surface-bond AuNPs may alter the effective aspect ratio or overall morphology of the nanostructure and influence colloidal stability.^20^ In addition, AuNPs attached to the origami surface may detach during handling, purification, or biological incubation due to mechanical shear forces, or nuclease-mediated degradation of the DNA linkages connecting the AuNP to the origami, particularly when these linkages are externally exposed. Consequently, AuNP detachment can lead to misidentification of barcodes, thereby compromising the validity of data interpretation.

In this work we develop a class of DNA origami barcodes, in which the AuNPs that form the barcode pattern are not exposed on the surface, but are wrapped inside tubular DNA origamis, which protects the linkages connecting the AuNPs to the origami from mechanical shear forces. The surface chemistry and aspect ratios of all barcodes are therefore identical, removing them as a potential bias in biologically relevant studies.

## Results and Discussion

### Barrel Design

In our first design, we modified a DNA barrel architecture from ref. ^21^ with an inner diameter of approximately 28 nm and a height of 22 nm (**Figure 1A**). Within the interior of the barrel, we incorporated four ss overhangs or “catcher” strands per site, that were designed to hybridize with complementary sequences on oligonucleotide-functionalized AuNPs (**Figure S1)**. In a separate reaction, 5 nm AuNPs were densely functionalized with thiolated oligonucleotides that are complementary to the capture strands at the inside of the origami.

**Figure 1.**
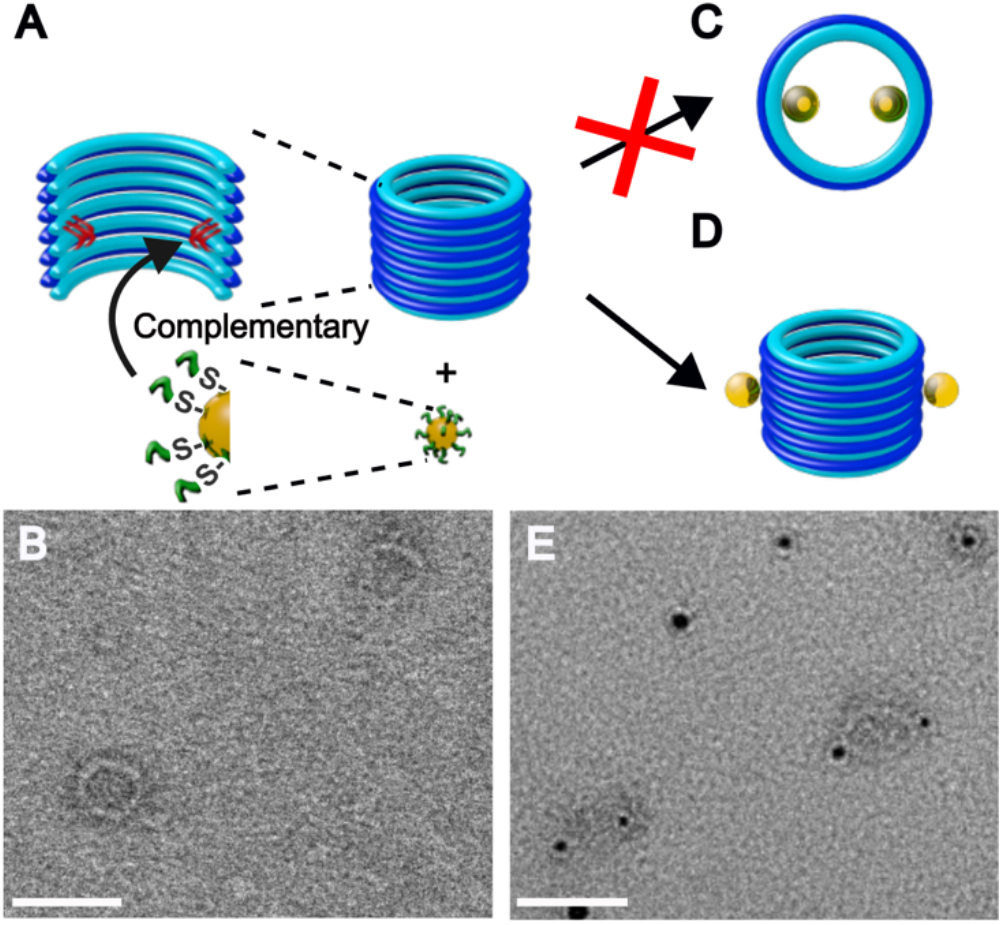
Unsuccessful DNA barrel design. (**A**) Schematic representation of the DNA barrel structure and AuNPs functionalized with complementary ssDNA. (**B**) Representative TEM micrograph of the barrel origami structure. (**C**) Following AuNP addition, inner encapsulation of AuNPs was not observed. (**D**) Instead, AuNP attachment occurred on the external surface of the barrel. (**E**) Representative TEM micrograph showing externally functionalized barrel structures. Scale bars: 50 nm.

This was followed by the incubation of the barrel origami structures with the AuNPs, to allow hybridization-mediated attachment. Agarose gel electrophoresis confirmed successful assembly (**Figure S2)**, and TEM micrographs revealed well-formed, circular DNA barrel structures (**Figure 1B**). However, most AuNPs were not located in the internal cavity as intended (**Figure 1C**) but were instead found on the external surface of the barrel (**Figure 1D** and **E**) or were missing. A possible explanation is that the wall of our modified design is only two helices thick, instead of three in the original structure.^21^ This reduction in wall thickness may have enabled a reorientation of the complementary catcher probe extensions and threading toward the exterior surface, resulting in hybridization of AuNPs to the outer wall rather than to the inside cavity. A similar observation has previously been reported and subsequently supported by molecular dynamics simulations.^22^ Moreover, the DNA modifications on the surface of AuNPs, together with the associated hydration shell consisting of tightly bound water molecules and ions, increases the effective hydrodynamic radius approximately two-to threefold. As a result, diffusion of these bulky particles into the centers of the barrels becomes entropically improbable, further reducing the yield of AuNP attachment within the cavity.

### Falafel-Wrap Design

For this reason, we abandoned the barrel design and developed a “falafel-wrap” inspired approach, in which AuNPs, *i*.*e*., “falafel balls”, are first attached to flat DNA origami sheets, *i*.*e*., the “flatbread”, and subsequently wrapped into tubular structures. To implement this approach, we modified the flat rectangular origami (FRO) originally reported by Rothemund,^8^ with overhangs on both faces of the structure. Initially, FRO structures were designed to incorporate three AuNP catcher strands at two distinct locations to promote tubular formation around two AuNPs (**Figure 2A**) with a diameter of 5, 10 or 20 nm. To induce closure into a tubular conformation, the two longitudinal edges intended for joining were designed to include four ss regions on each side of the scaffold. Eight complementary locking strands were subsequently added to bridge these regions and connect the opposing edges, stabilizing the closed tube configuration using a previously reported edge-locking strategy (**Figure S3**).^23^

**Figure 2.**
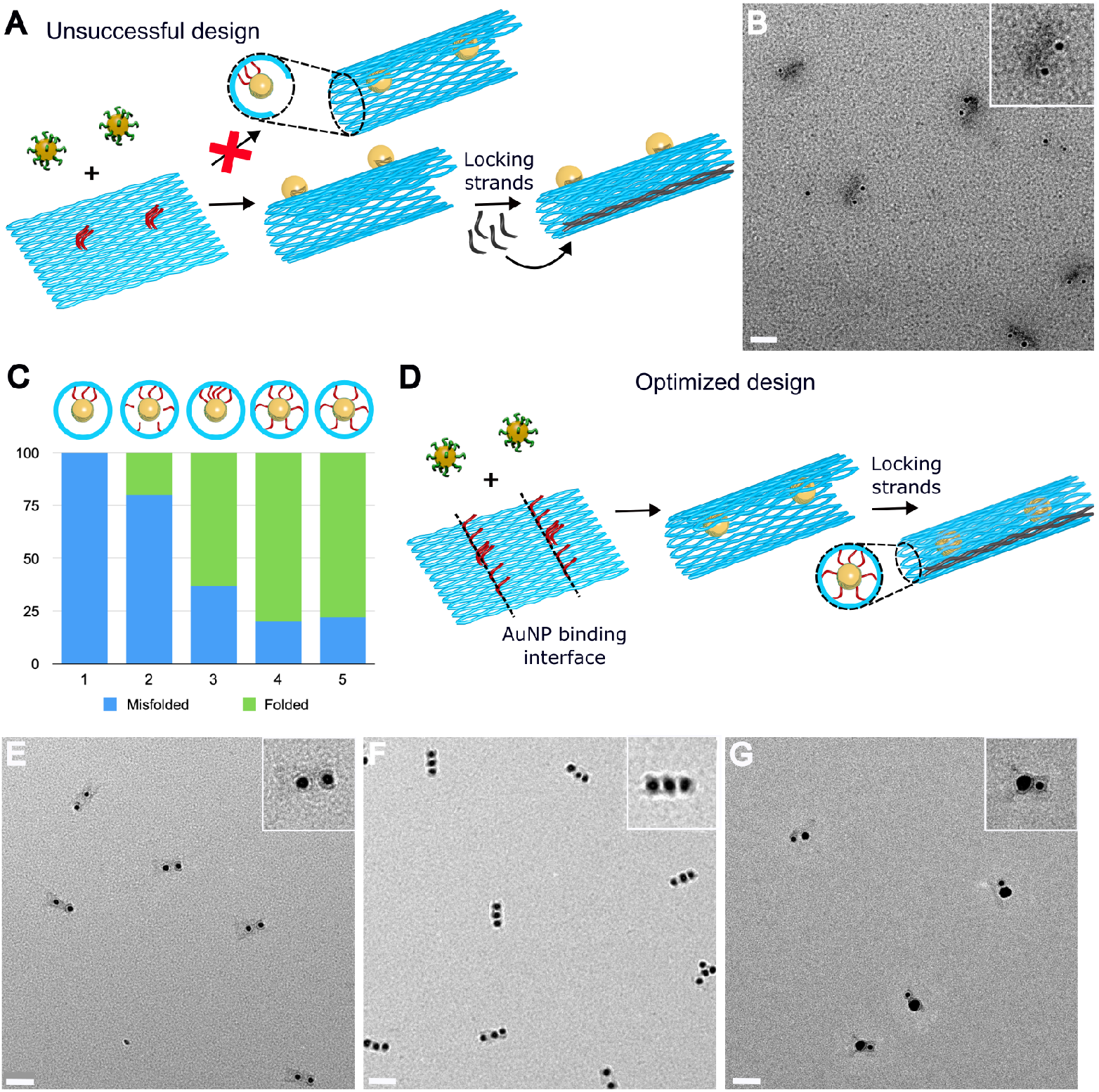
Falafel-wrap inspired design and optimization of handle position. (**A**) Schematic representation of the first design incorporating three ssDNA catcher strands per AuNP hybridization site, in which two 5 nm AuNPs end up on the outside. (**B**) Representative TEM micrograph of resulting origami structures obtained with this design. (**C**) Quantitative analysis showing the fraction of misfolded versus correctly folded structures for five different placements of additional AuNP catcher strands as drawn in the schematic cross section. (**D**) The optimized binding site design includes three central catcher strands and two longer peripheral. (**E-G**) Representative TEM micrographs of successfully folded structures encapsulating two 10 nm sized AuNPs (**E**), three 10 nm sized AuNPs (**F**), and 10 and 20 nm sized AuNPs of (**G**). Scale bars: 50 nm.

After barcode assembly and purification by agarose gel electrophoresis (**Figure S4**), TEM showed tubular structures in which the AuNPs were localized on the exterior surface rather than being encapsulated (**Figure 2A, B**). We hypothesized that the bulky, oligonucleotide-modified AuNP created an entropic brush effect, in a similar fashion as single-stranded overhangs in “hairygami” designs,^24^ inducing curvature in the wrong direction.

### AuNP-guided wrapping

To address this issue, we explored how different arrangements of AuNP capture strands on the FRO influenced the yield of correctly wrapped tubular structures with internally localized AuNPs. We hypothesized that the cooperative hybridization of capture strands distributed across the full width of the origami to AuNPs would generate curvature in the intended orientation where AuNPs end up on the inside of the tube. For that, we investigated four capture strand configurations: **(2)** three central catcher strands flanked by four additional handles, two on each side, distributed along the binding interface, to span approximately the AuNP diameter; **(3)** three central catcher strands with two short, 5 bp, peripheral handles on each side of the axis; **(4)** three central handles with two longer peripheral handles on each side of the axis; and **(5)** five equally spaced handles along the longitudinal axis of the FRO (**Figure 2D**). For configuration **(5)**, a ten-fold excess of AuNPs, as reported in the literature,^25^ led to multiple AuNPs binding per site, therefore, the incubation conditions were optimized to a three-fold AuNP excess to achieve controlled wrapping of the FRO around the AuNPs. Among the tested configurations, **(4)** exhibited the highest encapsulation efficiency. This design enabled correct inward localization of AuNPs across multiple barcode formats, including structures containing two equally sized AuNPs (**Figure 2E**), three equally sized AuNPs (**Figure 2F**), and two differently sized AuNPs (**Figure 2G**), with yields ranging from 60% to 80% of the assembled structures, as confirmed by TEM analysis.

### Functionalization of outer surface for cell targeting

We next explored the functionalization of the outer surface of the FRO for future applications in cell targeting, while maintaining the correct curvature where AuNPs are wrapped on the inside. For this, we explored a range of curvature-generating designs through coarse-grained oxDNA simulations, which are an excellent predictor of mechanical properties of DNA origami structures.^26,27^ The Šulc lab demonstrated that multiple single-stranded overhangs on one face of a FRO generates convex curvature through steric and electrostatic repulsion.^24^ Inspired by this, we varied overhang placement on one or both faces of the origami (**Figure 3**). Alternatively, we changed the number of base pairs between crossovers, which also induces curvature. Since full-scale simulations of large origami structures are computationally more expensive, we designed a smaller model version of the FRO to accelerate the simulation (**Figure 3** and **Figure S5**). Outer overhangs were designed to outnumber inner overhangs, and were also made approximately twice as long as the inner strands to reflect the shorter sequences required for AuNP functionalization.

**Figure 3.**
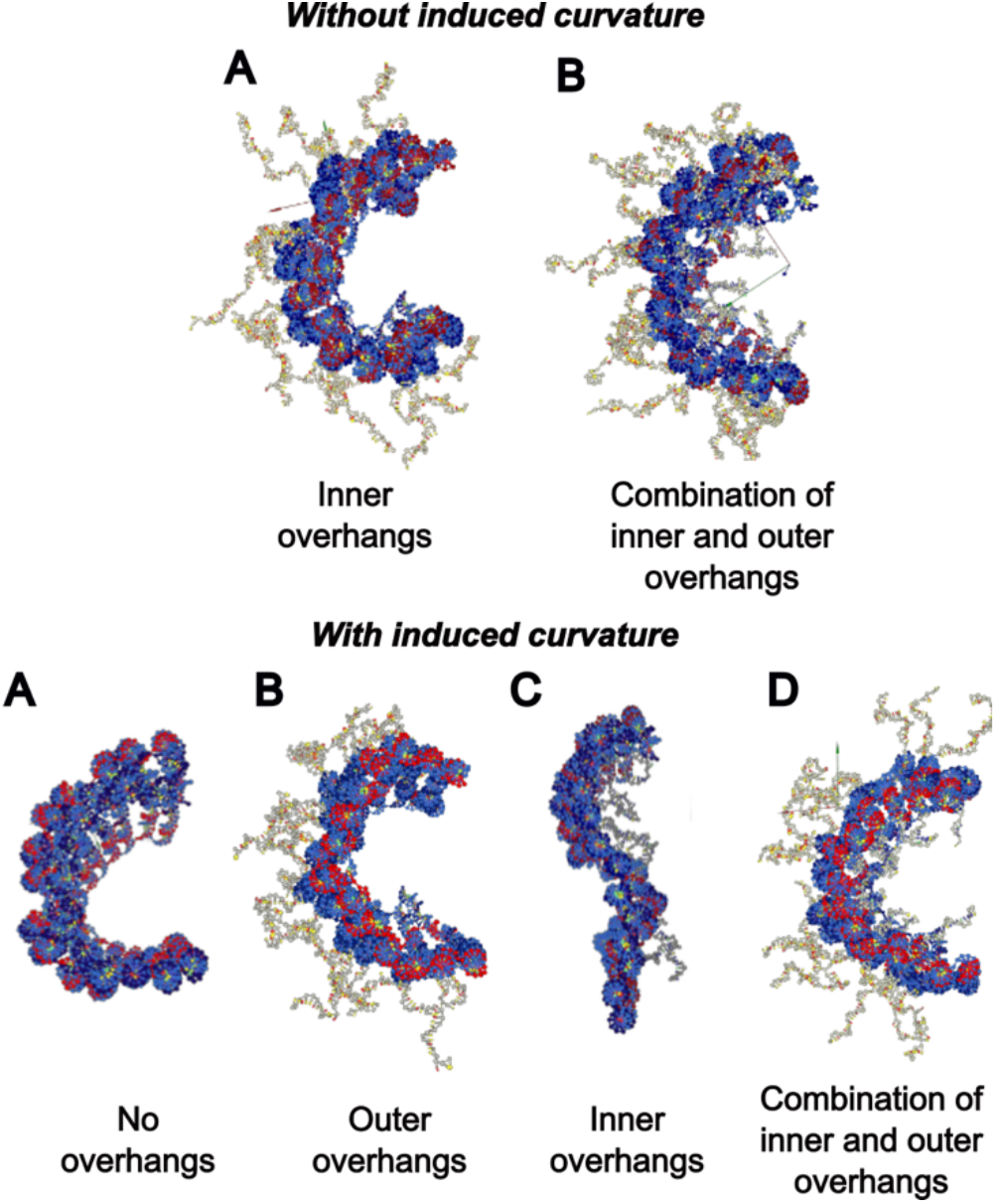
Coarse-grained oxDNA simulations of a smaller FRO model: without induced curvature. (**A**) inner overhangs; and (**B**) a combination of short inner and longer outer overhangs, and **with induced curvature** (**A**) no overhangs; (**B**) outer overhangs; (**C**) inner overhangs; and (**D**) a combination of short inner and longer outer overhangs.

Our simulation results confirm that structures designed with or without induced curvature, but containing overhangs on both surfaces, induced curvature toward the inner side (**Figure 3D**). This behavior arises because the steric and electrostatic repulsion generated by the more numerous and longer outer overhangs outweighs that from the inner overhangs, thereby directing the bending inward. This effect was further supported by the slight curvature observed when only inner overhangs were present and no pre-designed crossover shifts were introduced (**Figure 3C** Without induced curvature). Based on these findings, we selected the FRO design without additional pre-programmed crossover-induced curvature (**Figure S5A**), as the presence of outer overhangs alone was sufficient to promote favorable inward bending while minimizing unintended distortion. Following these simulations studies, we proceeded with experimental validation.

### Figure 3 Figure 3 Aptamer- and fluorophore-decorated barcodes

To evaluate additional functionalities, all accessible outer nick sites of the FRO origami were modified to accommodate either aptamers alone, a combination of aptamers and overhangs for subsequent hybridization-based attachment of fluorophores, or inner overhangs for fluorophore incorporation on the inner face (**Figure 4**). In these designs, up to sixteen binding sites were incorporated for an oligonucleotide carrying fluorescent dye cyanine 5 (Cy5). As a model targeting ligand, we selected the G-quadruplex-forming aptamer AS1411 due to its well established affinity for nucleolin receptors, which are overexpressed on the surface of many cancer cells, enabling selective cell targeting.^28,29^ In addition, the incorporation of fluorophores will provide an orthogonal optical readout, enabling fluorescence-based detection, tracking, and validation of barcode binding and cellular interactions in conjunction with TEM-based structural decoding.

**Figure 4.**
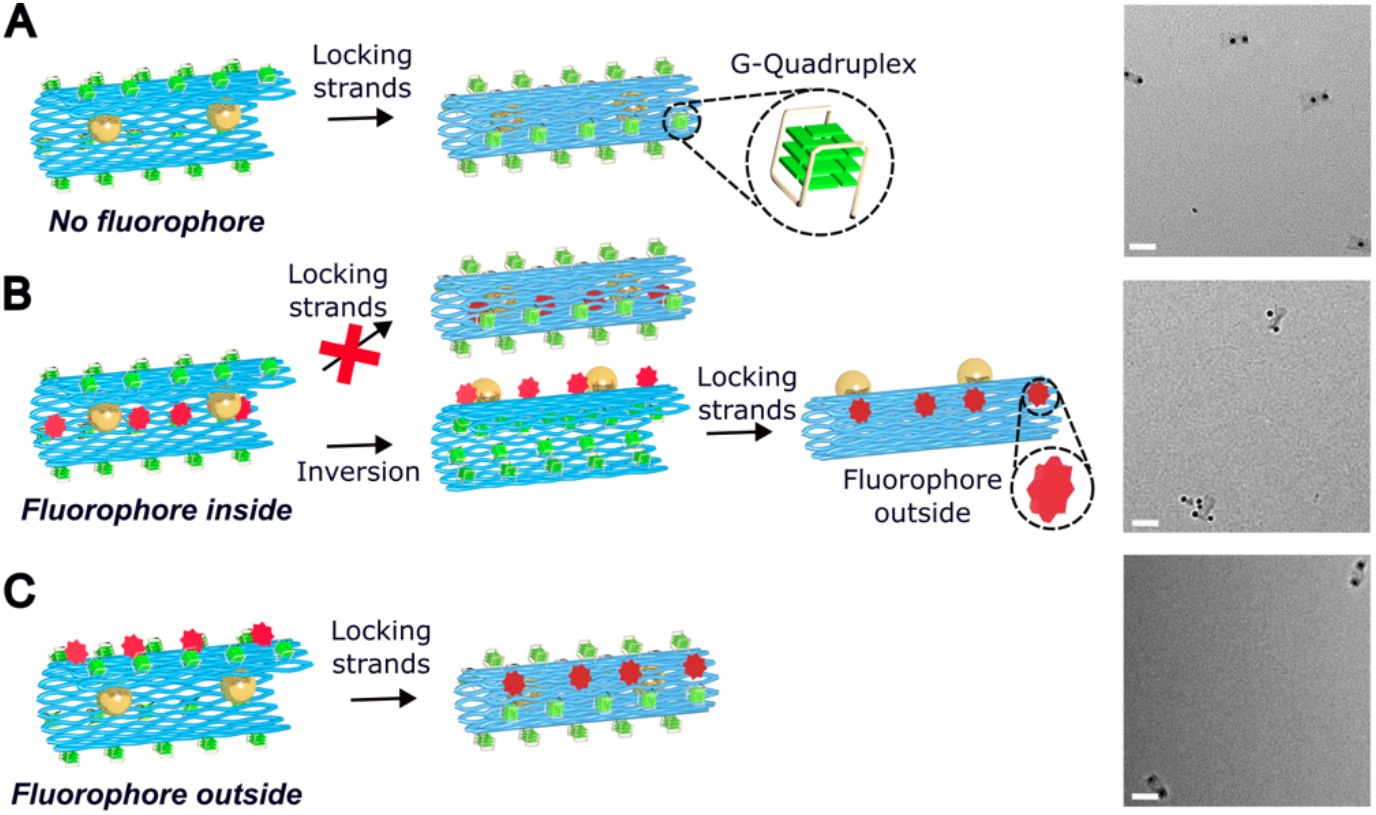
Functionalization strategies for aptamer- and -fluorophore-modified barcodes. (**A**) Design incorporating aptamers on the outer surface without fluorophores, and representative TEM micrograph; (**B**) Design combining outer-surface aptamers with inner-surface fluorophores, which resulted in inversion of the inner face to the exterior of the barcode, and the representative TEM micrograph; (**C**) Design incorporating both aptamers and fluorophores on the outer surface, and the representative TEM micrograph. Scale bars: 50 nm.

All barcodes were assembled using 10 nm AuNPs and these constructs were prepared following the same assembly protocol as described in **Figure 2**, with the addition of Cy5 during the AuNP incubation set to enable simultaneous hybridization (**Figure S6**). TEM analysis showed that all designs yielded the intended structures, except for the configuration incorporating Cy5 on the interior face of the FRO origami (**Figure 4B**). In this case, TEM micrographs revealed that the AuNPs were predominantly localized on the exterior of the barcode rather than being encapsulated. We hypothesize that the presence of sixteen Cy5 dyes on the inner surface generated increased steric and electrostatic repulsion, which overrode the inward curvature induced by AuNP binding and external aptamer functionalization.

### Barcode Dimerization

To further expand the diversity of possible barcodes, we designed two distinct monomer barcodes A and B, each displaying six external handles of 10 base pairs (bp) in length (**Figure 5A**). The handle sequences were designed to be mutually complementary between the two monomers, thereby preventing homodimer formation and enhancing the programmability for the generation of diverse dimer barcodes. Each barcode monomer encapsulated two 10 nm AuNPs. Dimerization was confirmed by agarose gel electrophoresis (**Figure S7**).

**Figure 5.**
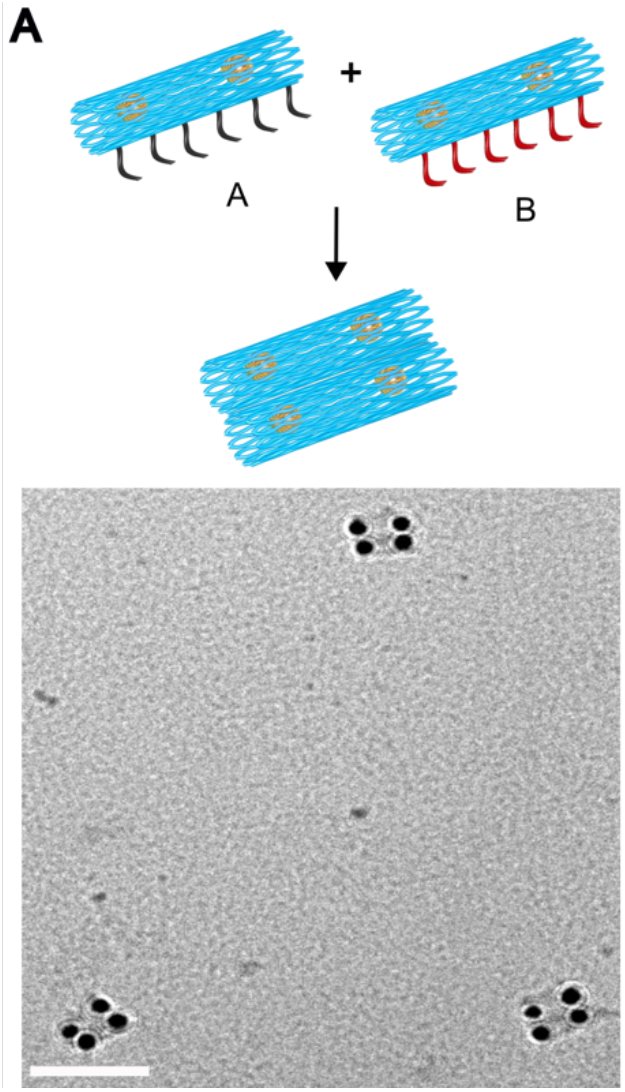
Barcode dimer assembly. (**A**) Dimer formation from two complementary monomer barcodes A and B. (**B**) Representative TEM micrograph of dimers. Scale bar: 100 nm.

We observed that incorporating short thymine spacers between the handles improved dimerization efficiency, likely by introducing additional conformational flexibility to accommodate thermal fluctuations and local structural deformations.^30^ Optimization revealed that incorporating 5 thymine residues resulted in the highest dimerization efficiency, yielding approximately 70%, as observed by TEM analysis (**Figure 5B**).

## Conclusion and Outlook

There is a growing demand for multiplexed detection and identification of various biomarkers, including nucleic acids and proteins, to enable efficient sample processing and high-throughput analysis. The use of DNA origami-based barcodes has emerged as a promising strategy to address this need, owning to their nanoscale addressability, structural programmability, and compatibility with high-resolution imaging techniques.^1–7^ In particular, the integration of AuNPs into DNA origami barcodes enables encoding of either the origami geometry or the identity of a specific biological target directly to the nanoparticle pattern, while allowing straightforward decoding by TEM imaging.^1,2^ A main limitation associated with the use of AuNPs is nanoparticle detachment, which can lead to misidentification or decoding errors in barcode analysis. Here, we demonstrate the controlled encapsulation of AuNPs within 3D origami structures by engineering inward curvature and increasing AuNP binding density at defined sites. We further show successful encapsulation of varying numbers of AuNPs, as well as the formation of dimeric assemblies to expand multiplexing capabilities. We further show successful encapsulation of varying numbers of AuNPs, as well as the functionalization of these barcodes with G-quadruplex-forming aptamers and fluorophores, enabling targeted cell recognition and providing an orthogonal fluorescence-based readout for tracking and validation of barcode-cell interactions. In addition, we show the formation of dimeric barcode assemblies to expand multiplexing capabilities. By confining AuNPs within the DNA nanostructures, nanoparticle loss is minimized and structural fidelity is preserved, providing a reliable barcode decoding. Together, these advances highlight the potential of AuNP DNA origami architectures for high-resolution, multiplexed biomolecular labeling and represent a significant contribution to the development of programable nanoscale diagnostic tools.

## Associated content

Supporting information contains experimental methods, caDNAno designs, and electrophoresis data.

## Author Information

### Author contributions

D.P.N.G. and T.L.S. conceived the study; S.Y. performed experiments; T.S. and S.Y. performed coarse-grained molecular dynamics simulations; D.P.N.G. and T.L.S. acquired funding; D.P.N.G. and T.L.S. supervised the study; D.P.N.G., T.L.S. and S.Y. wrote the initial and final draft; all authors revised and expanded the manuscript.

### Notes

The authors declare no competing financial interest.

## Acknowledgements

We wish to acknowledge Dr. P.S. Prakash for her valuable help and suggestions regarding AuPN functionalization of the barcodes and meaning full discussions.

## Materials and methods

### Barrel Design

The barrel design was adapted from the architecture reported by Wickham *et al*.,^1^ which features regularly spaced nick points at diagonal intervals of 8 bp. To increase the internal volume for AuNP accommodation, the miniscaffold present in the original design was removed, and the inner handles were extended from the inner layer for AuNP hybridization. Nick site placement for handle incorporation was designed using caDNAno software.^2^ The scaffold strand (10 nM, P7249, Tilibit) was mixed with a 10-fold excess of staple strands (Eurofins) in 1x Tris-Edetate buffer (TE, 5 mM Tris, 1 mM EDTA, Sigma-Aldrich) supplemented with 10 mM MgCl_2_. Barrel structures were assembled in a thermocycler (Eppendorf) by heating to 80 °C, cooling to 65 °C over 15 min, and subsequently maintaining the temperature at 47 °C for 18 h.

### FRO Design

The FRO design was adapted from the flat rectangular origami reported by Rothermund,^3^ with modifications to introduce additional nick sites on the face opposite to the original nicks, thereby enabling handle extensions from both sides of the structure. Although the adapted design aimed to introduce an equal number of nicks on both faces, placing nicks near the central seam produces staples that are too short, leading to misfolding in that region. As a result, the final design contained an asymmetric distribution of handles. The face with a higher density of handles was designated as the outer surface for aptamer and/or fluorophore functionalization, while the face with fewer handles was assigned as the inner surface for AuNP encapsulation. The DNA nanostructures were assembled by thermal annealing using a temperature ramp from 80 to 55 °C at a rate of 1 °C/min, followed by cooling from 55 to 25 °C over 1 h, in 1x TE buffer supplemented with 12.5 mM MgCl_2_.

### Purification of DNA origami samples

After thermal annealing, the structures were purified using 100 kDa molecular weight cutoff (MWCO) filters (Amicon Ultra, Millipore) by centrifugation at 10,000 rcf for 5 min per spin cycle. The spin buffer consisted of 1x TE buffer supplemented with 5 mM MgCl_2_ and 300 mM NaCl. The number of spin cycles was adjusted proportionally to the sample volume and continued until no excess staple smear was detected by gel electrophoresis. The concentration of the purified origami was then determined using a NanoDrop spectrophotometer (Thermo Fisher Scientific) at 260 nm.

### Functionalization of AuNPs with ssDNA

AuNP functionalization with ssDNA was performed following a modified procedure based on Prakash *et al*.^4^ In brief, the citrate layer on AuNPs (5, 10, or 20 nm, Sigma-Aldrich) was first exchanged with a phosphine ligand by adding 5 mg of bis(*p*-sulfonatophenyl)phenylphosphine dihydrate dipotassium salt (BSPP, Sigma-Aldrich) to 12.5 mL of the AuNP solution and incubating overnight. Subsequently, 5 M NaCl was added dropwise until a color change from red to blue-violet was observed. The mixture was then centrifuged at 1600 rcf for 30 min at 4 °C for all particle sizes except 5 nm AuNPs, which were centrifuged at 3000 rcf. The supernatant was discarded, and the pellet was resuspended in a solution containing equal volumes of 2.5 mM BSPP and methanol (MeOH, VWR). After a second centrifugation step, the AuNPs were resuspended in 1 mL of 2.5 mM BSPP solution. The absorbance of the AuNP suspension was measured at 525 nm, and the nanoparticle concentration was calculated using the excitation coefficient provided by the manufacturer. The functionalized AuNPs were stored at 4 °C in the dark until further use.

Thiol-modified ssDNA strands were first treated with 20 mM tris(2-carboxyethyl)phosphine hydrochloride (TCEP, Sigma-Aldrich) for 45 min to reduce disulfide bonds and generate free thiols. The reduced ssDNA was then added to the AUNP solution, briefly sonicated (10 s), and incubated in the dark for 20 min. An excess of ssDNA was used to ensure maximal surface coverage of the AuNPs (molar ratios: 1:5 for 5 nm, 1:150 for 10 nm, and 1:450 for 20 nm AuNPs). To promote efficient DNA absorption, a salt-aging process was carried out by gradually increasing the NaCl concentration to 1M (25, 50, 100, 200, 400, 700, and 1000 mM). After each addition, the samples were sonicated for 10 s and incubated for 20 min. The DNA-functionalized AuNPs were then gently agitated on a shaker (25 rpm) overnight. Excess ssDNA was removed by purification using pre-wetted 100 kDa MWCF, followed by at least 8 centrifugation cycles in 0.5x TE buffer at 10,000 rcf for 5 min per cycle. The concentration of the functionalized AuNPs was determined by measuring the absorbance at 525 nm.

### Barcode assembly

For barcode assembly, DNA origami samples were mixed with AuNP stock solutions to achieve a final concentration of 10 nM origami and a 3-4-fold excess of AuNPs per binding site, unless otherwise specified. Samples were either incubated for 2 h, followed by the addition of locking strands and overnight incubation at room temperature, or subjected to a controlled temperature ramp from 40 to 20 °C overnight.

### Dimer barcode assembly

For dimer assembly, barcodes were prepared using the protocol described above, with the inclusion of 10 bp complementary external handles for dimer formation. The two complementary barcodes were then incubated in an equimolar ratio overnight using a hybridization ramp from 40 to 20 °C.

### Barcode purification

Excess AuNPs were removed from the barcodes by agarose gel purification. Bands corresponding to the assembled barcodes were excised and gently compressed between two glass slides covered with parafilm to extract the barcode samples. The recovered material was then used for TEM characterization.

### Agarose gel electrophoresis

Barcodes were analyzed and purified by agarose gel electrophoresis using 1% agarose (Fisher Bioreagents) cast in 0.5x TBE buffer supplemented with 5 mM MgCl_2_. Gels were pre-stained with 1x SYBR Safe DNA gel stain (Invitrogen) and run at 70 V for 2 h. samples (10 µL) were mixed with 1x Orange DNA loading dye (6x stock, Thermo Scientific) prior loading. The same buffer was used as the running buffer, and electrophoresis was performed using a Biorad Sub-Cell GT horizontal electrophoresis system equipped with a Biorad PowerPac Basic power supply, at room temperature. Gels were subsequently imaged using a fluorescence scanner.

### TEM characterization

Samples (10 µL) were deposited onto plasma-cleaned, carbon-coated grids (400 mesh copper, carbon on Formvar, Ted Pella, Inc.) and incubated for 5-7 min. Excess liquid was removed using filter paper, followed by staining with 2% uranyl acetate (Ted Pella, Inc.) for 1 min. The staining solution was pre-activated with 0.1 M sodium hydroxide (NaOH, VWR) for 30 s. Excess stain was removed by two washes with 5 µL Milli-Q water, and the grids were allowed to air-dry. Imaging was performed using an FEI Tecnai G2 transmission electron microscope, and images were analyzed using Fiji (ImageJ) software.

### oxDNA simulations

Curvature studies of the FRO were performed using coarse-grained oxDNA simulations on a reduced model approximately 100 bp in width. Insertions and deletions were introduced into the design to evaluate their effect on curvature. Simulations were carried out using oxView online platform following established protocols.^5^ Briefly, structures were generated using the “create from base pairs” function and submitted to the Nanobase server for simulation. Initial CPU-based simulations were performed at 20 °C using default parameters. Each simulation was run until the potential energy reached approximately -1.4 or until convergence was observed with no further significant changes in energy.

### Assembly yield and structural integrity analysis

Structures exhibiting the correct barcode design were counted, with at least 50 structures analyzed per sample. The percentage was calculated relative to the total number of counted structures, including both correctly folded and misfolded assemblies.

---

**Figure S1.**
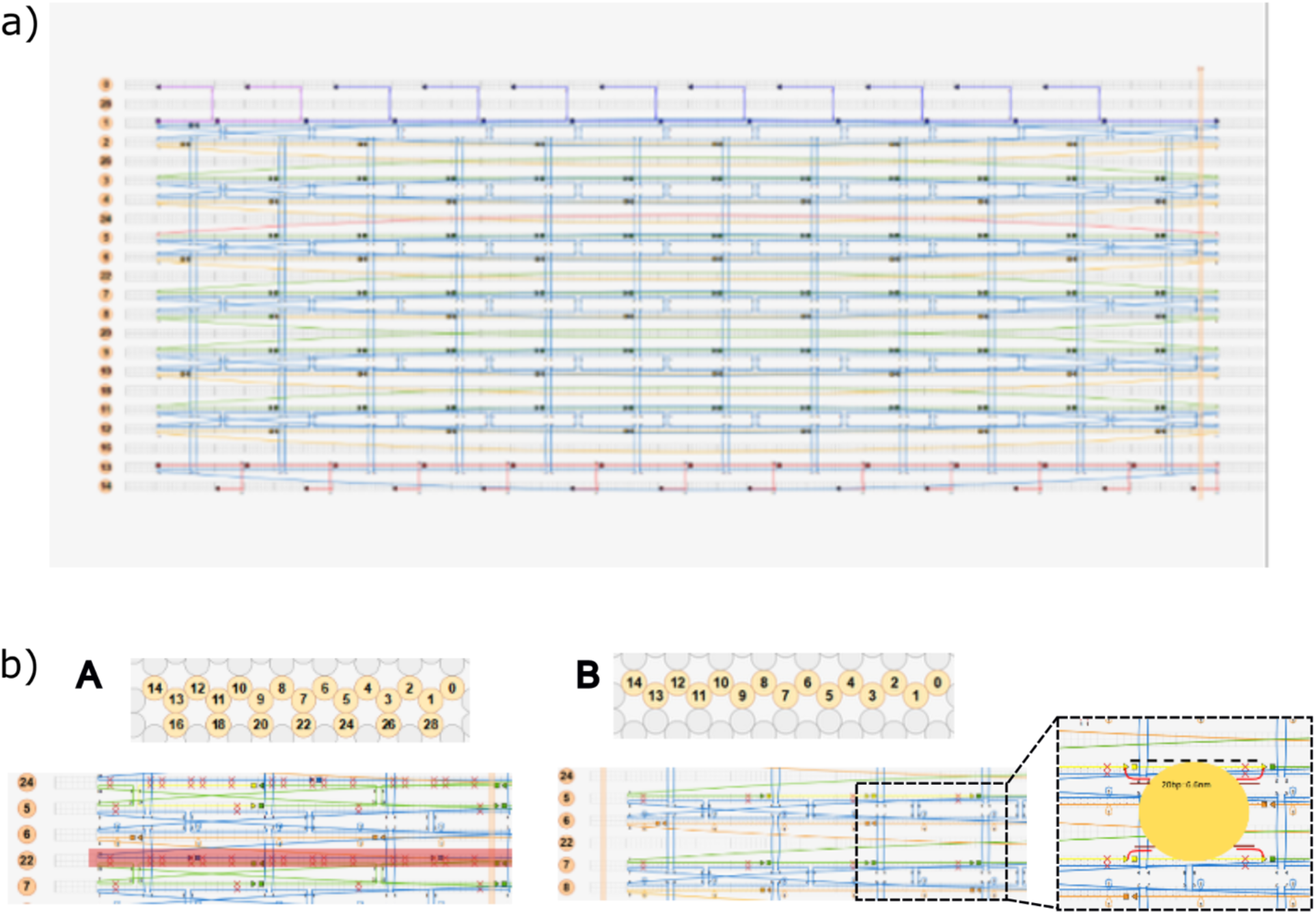
Barrel design overview. **a)** caDNAno design of the modified barrel structure, in which the inner miniscaffold has been removed, and selected nick sites (green) have been reoriented toward the barrel cavity. Red circles indicate the positions of ss overhangs for AuNP attachment via hybridization. Dashed boxes denote individual binding sites, each containing four overhangs designed to capture a single AuNP. Staples at the top, bottom along with their corresponding nick sites (orange), were designated for outer-surface aptamer functionalization. **b)** Comparison of the barrel design by Wickham et al., with the modified one. (**A**) Original design. (**B**) Modified design, with detailed view of the inner overhand location for AuNP attachment.

**Figure S2.**
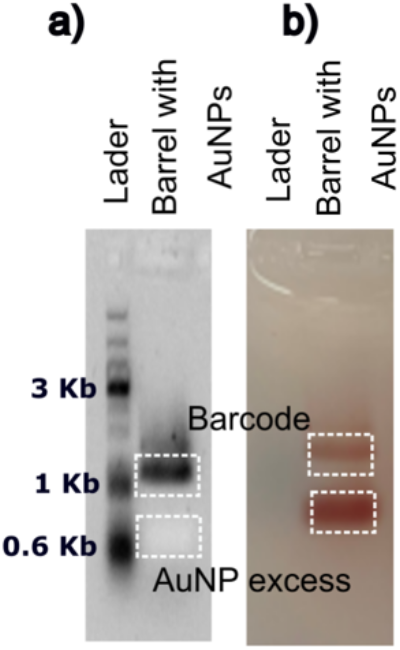
Agarose gel of the barrel structure following incubation with excess 5 nm AuNPs. **a)** Fluorescence image of the gel stained with SYBR Gold. **b)** Corresponding color photograph of the gel.

**Figure S3.**
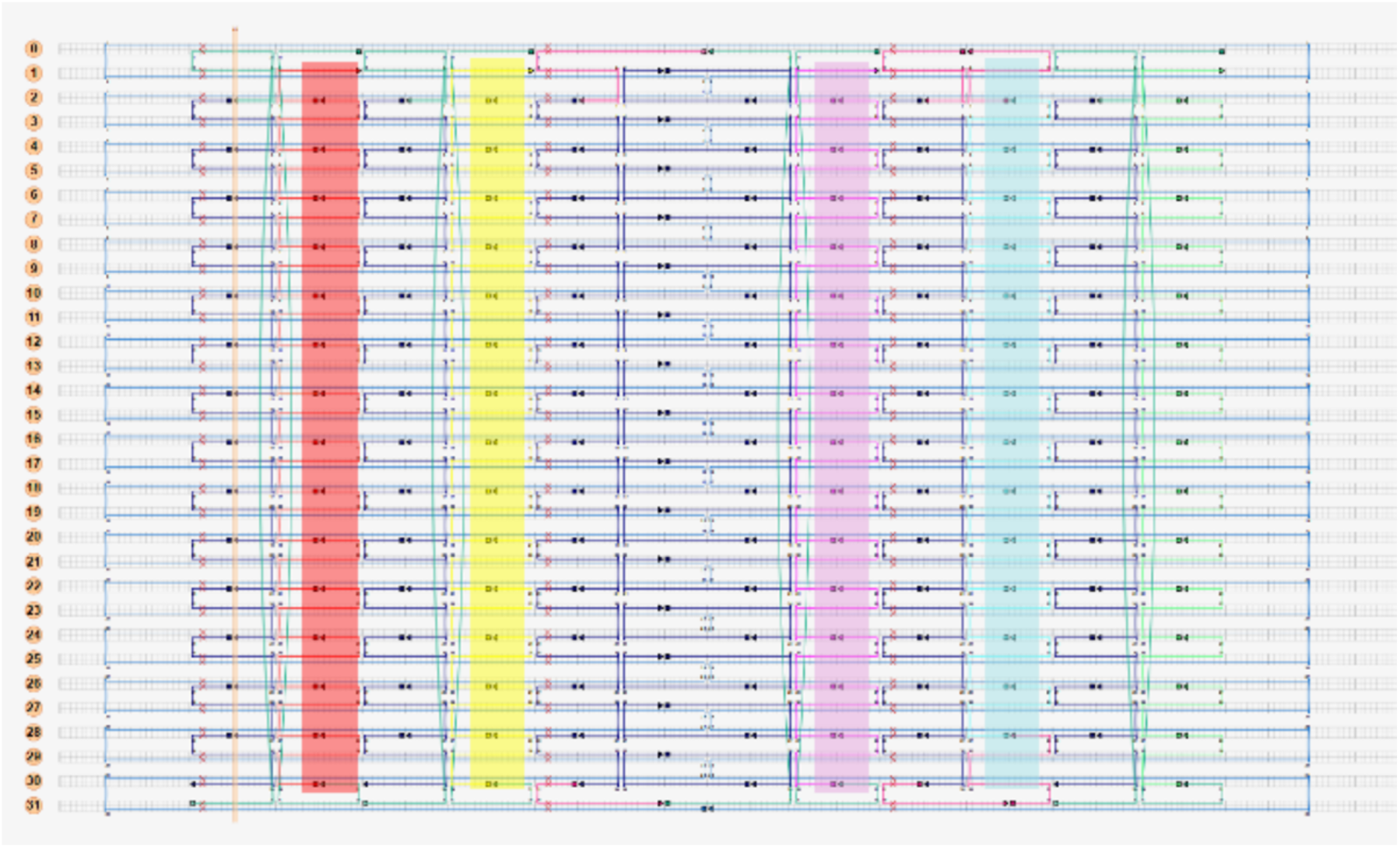
Design overview of the FRO. caDNAno design of the FRO structure highlighting all nick sites. Outer overhang extensions are shown in dark blue, while colored rectangles indicate the potential AuNP hybridization sites, with each color corresponding to a single AuNP position. Sites that will hybridize with the locking strands are shown in green.

**Figure S4.**
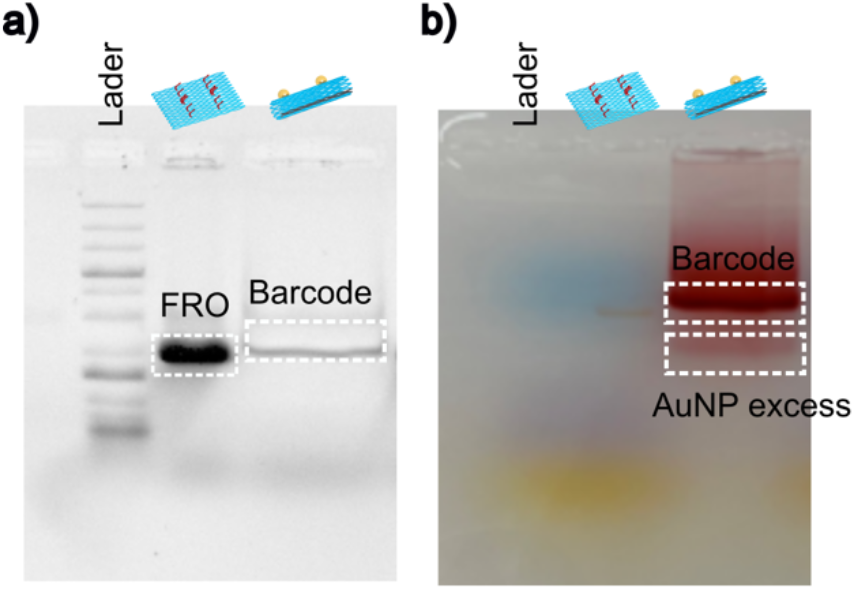
Agarose gel of FRO structure before and after incubation with excess 10 nm AuNPs. **a)** Fluorescence image of the gel stained with SYBR Gold. **b)** Corresponding color photograph of the gel.

**Figure S5.**
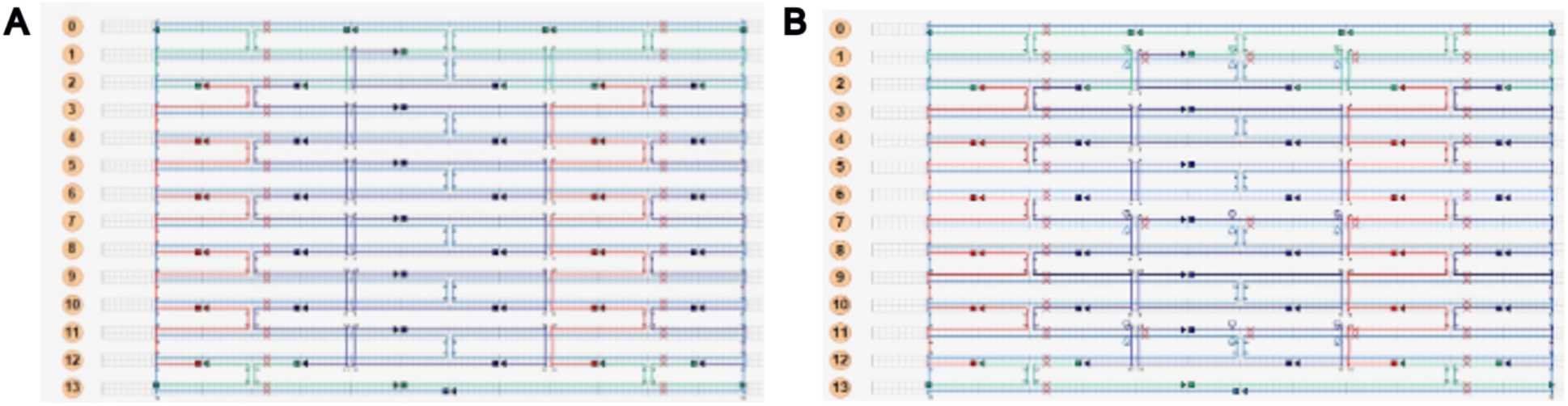
caDNAno design of the reduced FRO model used for oxDNA simulations. (**A**) FRO without pre-designed curvature. (**B**) FRO incorporating left-handed crossover shifts to induce curvature.

**Figure S6.**
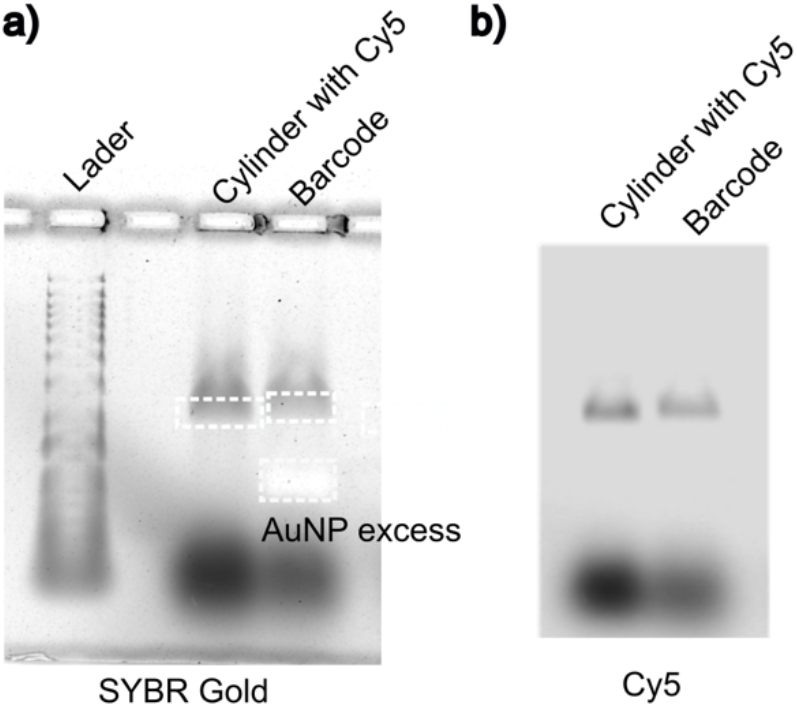
Agarose gel analysis comparing FRO cylinders functionalized with G-quadruplex forming aptamers and Cy5 on the outer surface to of barcodes functionalized with G-quadruplex forming aptamers and fluorophores on the outer surface. a) Fluorescence image of the gel stained with SYBR Gold. b) Fluorescence image acquired in the Cy5 channel.

**Figure S7.**
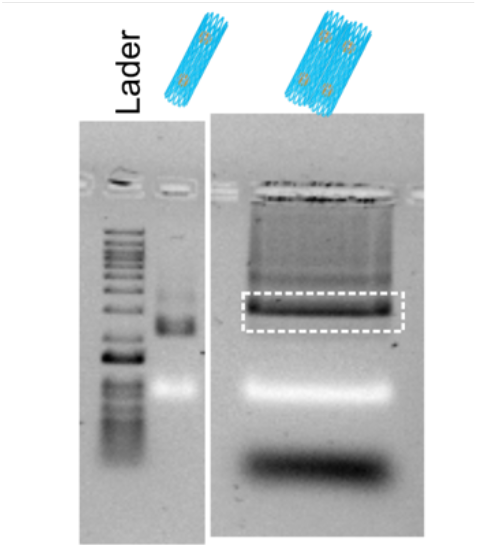
Agarose gel of the barcode dimer after assembly.

